# Host-strain compatibility influences transcriptional responses in *Mycobacterium tuberculosis* infections

**DOI:** 10.1101/2024.05.21.595142

**Authors:** Paula Ruiz-Rodriguez, Marta Caballer-Gual, Guillem Santamaria, Hellen Hiza, Mireia Coscolla

**Affiliations:** Institute for Integrative Systems Biology (I^2^SysBio), University of Valencia-CSIC, FISABIO Joint Research Unit Infection and Public Health, Valencia, Spain; Luxembourg Centre for Systems Biomedicine (LCSB), University of Luxembourg, Esch-Belval Esch-sur-Alzette, Luxembourg

**Keywords:** *Mycobacterium tuberculosis*, host-pathogen interactions, RNA-seq, transcriptomics, infection ratio

## Abstract

Tuberculosis, caused by *Mycobacterium tuberculosis*, is a leading cause of infectious mortality and affects humans and other mammals. Understanding the drivers of its host-specificity shapes the landscape of infectivity, which could potentially inform strategies for tuberculosis management. We hypothesise that host-strain compatibility influences infection outcome and we aim to reveal bacteria and host specific transcriptional responses during *in vitro* infections.

We infected human and bovine cell lines with two human-associated *M. tuberculosis* strains from lineages 5 and 6, as well as two animal-associated strains (*Mycobacterium bovis* and the Chimpanzee bacillus), and quantified infection ratios, cell death, and transcriptional responses. By integrating those data from different compatibility combinations, we identified global transcription profiles associated with strain-host compatibility.

Our results show that the most compatible host-strain combinations had higher infection rates, and different transcriptional patterns than low-compatibility infections. Both hosts had similar transcriptional responses to their most compatible strains, upregulating pathways related with increased cell proliferation. Host proliferation could potentially favour bacterial growth, explaining the success of the strain in its corresponding host. Conversely, both hosts responded to non-associated strains with defence related transcriptional patterns, among other pathways, supporting their lower success in the non-associated host. Finally, we revealed bacteria and host-specific expression patterns in molecules implicated in immune response and inflammation through the eicosanoid pathway.

In conclusion, we confirmed that bacteria-host compatibility determines common transcriptional responses, but also specific responses that depend on the infecting strain. This study enhances the understanding of host-specific adaptation mechanisms in *M. tuberculosis*.

## Introduction

Pathogen specificity, the phenomenon where pathogens exhibit a preference for infecting certain hosts over others, is a cornerstone concept in understanding the dynamics of infectious diseases. There are ecological, epidemiological, and biological drivers of such specificity, where hosts provide more permissive conditions to support the pathogen’s life cycle. Biologically, this specificity can also result from compatibility between interacting molecules resulting in increased efficiency entering the cell, resistance to the immune system, or replication within the host [1,2]. The implications of host specificity extend beyond basic science, affecting disease control strategies, vaccine development, and the management of emerging infectious diseases, especially within the context of One Health.

Pathogens displaying host specificity, such as influenza A, exhibit discrete molecular changes that are responsible for the host specificity [3]. Coronaviruses also display specificity for different hosts, and research into cross-species transmission events improves understanding of potential zoonotic spillover events [4,5]. However, in other pathogens like *Brucellaceae*, where different species infect different hosts, pinpointing such changes is challenging presumably because of the ancestral nature of host adaptations [6]. *Mycobacterium tuberculosis* also displays host specificity across different mammals [7], and the study of the molecular determinants of such specificity can potentially inform specific virulence determinants of bacteria-host combinations.

Tuberculosis ranks among the leading causes of death from a single infectious agent, and it is caused by members of *M. tuberculosis* [8]. *M. tuberculosis* includes human-associated *M. tuberculosis*, which consists of ten phylogenetic lineages (L1 to L10), and four animal-associated lineages: A1 to A4 [9–12]. Lineage A1 infects wild terrestrial animals in Africa and is the least studied among the animal-associated lineages despite being the closest relative to the human-associated Lineage 6 [13]. In contrast, Lineage A4, which includes *Mycobacterium bovis* and *Mycobacterium caprae,* is the most studied animal-associated clade and is mostly found in cattle and goats, respectively. Despite the preference of certain lineages for specific hosts, there is no absolute exclusivity in host tropism [14]. For example, wildlife reservoirs such as wild boars, badgers, or ferrets among others, can also harbour *M. bovis*, leading to spillback into cattle, which can, in turn, infect wildlife. *M. bovis* does occasionally infect humans, but human-to-human transmission of *M. bovis* is uncommon [15,16]. On the other hand, human-associated lineages are occasionally isolated from cattle and other animals due to contact with humans [9,13,17]. These infections are characterised by an attenuated virulence phenotype, with transmission back to humans being extremely uncommon [18,19].

Host jumps are concentrated in the clade encompassing all four animal-associated lineages as well as the human-associated Lineages (L) 5, 6, 9, and 10, which are among the least studied members of the species. Among these, L5, L6, and A4 have the most data available. Human-associated L5 and L6 are often referred to as *Mycobacterium africanum*, though they differ significantly. While L5 is associated with Ewe Ethnicity in West Africa [20], Lineage 6 is more diverse, shares a more recent common ancestor with the animal-associated clades, and it has been associated with HIV infection [21]. These factors suggest a potential difference in host specificity between L5 and L6, with L6 possibly having a more generalistic nature compared to L5, which has been associated with the Ewe ethnicity [20]. Conversely, Lineage A4 is more commonly associated with cattle, but the host range of other animal-associated lineages is not as clear due to the scarce data available. The lack of data is particularly significant for A1, including the Chimpanzee bacillus [22] which has been isolated only once despite being very closely related to L6.

It is not clear which are the molecular factors driving host specificity in *M. tuberculosis*. Phospholipase C (plc) genes are promising candidates, given the fact that they are prone to be lost in animal-associated lineages at different points during *M. tuberculosis* diversification [9,23]. Phospholipase C are proteins that are important in the virulence mechanisms of several pathogens such as *Listeria monocytogenes* [24] and *Pseudomonas aeruginosa* [25]. The role of these enzymes is less clear in *M. tuberculosis*, but some studies point to their involvement in virulence [25]. The *plc* genes in *M. tuberculosis* have been shown to block the eicosanoid signalling pathway, which produces a shift from apoptosis to necrosis [26]. In particular, they inhibit the synthesis of prostaglandin E2 (PGE2) and the expression of *PTGER2* and *PTGER4* [26]. These signalling molecules are crucial in mediating eicosanoid-derived innate immune response, which is critical for infection control [27].

Our goal is to advance the understanding of the molecular determinants of *M. tuberculosis* specificity using an *in vitro* infection system. We hypothesise that the Lineage 5 strain (L5) and the *M. bovis* (A4) strain will display a common pattern of high-virulence-related phenotypes in human and bovine cells, respectively, due to their higher compatibility. Conversely, Lineage 6 (L6) and the Chimpanzee bacillus (A1) will not exhibit such a clear pattern of virulence-related phenotypes in either host cell. To test this, we will examine whether the epidemiological compatibility between four different strains and their most common hosts can be partially mimicked by more successful infections in *in vitro* experiments. If so, we aim to identify the molecular mechanisms involved through expression analysis.

## Material and Methods

### Cell lines and *M. tuberculosis* strains

Human THP-1 and bovine macrophages (BoMac) were used for infections. Cell cultures were maintained in respective mediums before seeding and differentiation for each infection. THP-1 monocytes were cultured at 37°C with 5% CO_2_ to confluence in RPMI media (MEDIUM 1640 CE, Gibco, 21875034) supplemented with 10% FBS, then differentiated into macrophages using 15 ng/ml of phorbol 12-myristate 13-acetate (PMA) for 24h. BoMac were thawed and maintained at 37°C with 5% CO_2_ to confluence in IMDM media containing GlutaMax and HEPES (Gibco, 31980-022) supplemented with 10% FBS, 1x MEM Non-Essential Amino Acids (Gibco, 11140-035), 1x MEM Vitamins (Biochrom AG, K0373) and 50 μM β-Mercaptoethanol (Gibco, 31350-010). Macrophages were detached using trypsin-EDTA 25% (Gibco, R001100) and seeded in both 24- and 6-well cell culture-treated plates according to the intended procedure. All cells were rested for 24 hours before infection at 37°C with 5% CO_2_ to allow adherence of macrophages to the plates.

We infected with two strains from human-associated *M. tuberculosis* lineages: N1176 (L5) and N1202 (L6) [28] and two strains from animal-associated lineages: G1352 (*M. bovis*) [29], and N1894 (Chimpanzee bacillus) [22] (see Figure S1A and S1B).

### M. tuberculosis infections

All strains were grown at 37°C to mid-log phase (OD_600_ = 0.5 to 0.6) in Middlebrook 7H9 broth, enhanced with 10% (v/v) albumin/dextrose/catalase (ADC Middlebrook Enrichment – BD, 212352), 0.44% (w/v) sodium pyruvate, and 0.05% (v/v) Tween 20. Single cells from each strain were prepared by resuspending and washing the cultures in phosphate-buffered saline (PBS), resuspending in either supplemented RPMI or IMDM then followed by sonication at maximum speed for 2 minutes.

Rested 7·10^5^ THP-1 and 5·10^5^ BoMac macrophages in 6-well plates, and 5·10^5^ THP-1 and 1.8·10^5^ BoMac in 24-well plates were then infected with all strain-cell combinations at different multiplicity of infection (MOI) or left uninfected. MOIs of 10:1 and 20:1 were used in THP-1 and BoMac cells respectively. After 4 hours post-infection (hpi), extracellular bacteria were removed, and the media was replaced with RPMI or IMDM supplemented with 5 µg/µL gentamicin. After 1-day post-infection (dpi), the gentamicin-supplemented media was substituted with fresh RPMI or IMDM supplemented with 10% FBS. Infected cells were maintained for 6 days. Infections were done in replicates for each strain doing 26 infections in total.

### Infection ratio by microscopy

Optical microscopy analysis were performed from infected cells in 24 well plates, starting with 5·10^5^ THP-1 and 1.8·10^5^ BoMac cells per well. The infection ratio was determined by optical microscopy at 4 hpi, 1- and 3 dpi (see Figure S1C). Initially, THP-1 cells or BoMac cells were fixed with 3.7% formaldehyde. Subsequently, the infected cells were stained using the Ziehl-Neelsen technique. Cells were first stained with carbol fuchsin (Sigma-Aldrich, 108512), resulting in a pink stain on the infected cells followed by incubation in hydrochloric acid in ethanol (Sigma-Aldrich, 100327), leading to the decolorization of macrophages but not acid-fast cells, which retained their pink coloration. Finally, the decolorized macrophages were counter-stained with methylene blue (Sigma-Aldrich, 101287). We considered an infected cell as a macrophage containing at least one bacillus. To determine the infection ratio, both infected and uninfected cells were counted within a 44 mm^2^ area. The infection ratio was calculated by dividing the number of infected cells by the total number of cells in the observed area. We obtained data across 12 experiments with at least two technical replicates for each infection combination (Table S1).

### Cell death assessment by flow cytometry

To investigate macrophage cell death induced by different strains, we performed flow cytometry analysis at 2 and 6 dpi from infected 6-well plates starting with 7·10^5^ THP-1 and 5·10^5^ BoMac cells per well (see Figure S1C). The culture medium was removed from the wells, and the cells were detached with trypsin, and washed with PBS. The cells were then resuspended in 500 μL of PBS and then stained following the Fixable Viability Stain 450 kit manufacturer instructions (BD Biosciences, 562247). Infected and uninfected macrophages were stained with Fixable Viability Stain 450, diluted 1:10 in PBS, and incubated at 37°C for 5 minutes in the dark then washed with PBS. Subsequently, the samples were stained with 80 μL CellEvent Caspase 3/7 Green detection Reagent 2 µM in PBS with 5% FBS (Invitrogen, C10423) and then incubated for 30 minutes at 37°C. Stained cells were fixed with 80 μL of 7.4% formaldehyde for 10 minutes at room temperature. Finally, the supernatant was removed by centrifugation, and the cells were resuspended in 500 μL of PBS and then were read on a BD LSRFortessa the same day. FCS files obtained were exported and then analysed using FlowJo^TM^ Software v. 10.9. After obtaining cell viability percentages for each sample, we excluded controls identified as outliers using the interquartile range (IQR) method. We then computed the mean values of the controls for each experiment. Following this, the fold changes for the samples were calculated by dividing the sample values by the control means. We obtained data across 12 experimental setups with at least 2 technical replicates for each infection combination (Table S2). We removed values below 1 for apoptosis and necrosis, and we excluded values above 1 for live cells, if one category failed all the sample was removed. Additionally, outliers in the fold change detected with interquartile range (IQR) method were also removed. After filtering those low-quality data, we analysed data from 12 experiments with at least 2 technical replicates for each infection combination.

### RNA extraction and sequencing

RNA was extracted at 1- and 3 dpi (see Figure S1C) from infected cells in 6-well plates, starting with 7·10^5^ THP-1 and 5·10^5^ BoMac cells per well. We used 500 μL RNAzol per 5·10^5^ macrophages, followed by centrifugation for 20 min at 10000 rpm. The supernatant was removed for eukaryotic RNA sequencing, and the pellet was subjected to *M. tuberculosis* RNA extraction. Pellets from 6 wells were pulled and resuspended in 300 μL RNAzol and transferred to Lysing Matrix B containing 0.1 mm silica beads (MP biomedical) followed by vortexing for 3 minutes. Post-centrifugation, the supernatants containing *M. tuberculosis* RNA were transferred into a new tube. Thereafter, bacterial and eukaryotic RNAs were subjected to the RNA purification process. Isopropanol was added to induce RNA precipitation, followed by ethanol-washing steps to purify the RNA and remove impurities. Finally, the purified RNA from each sample was resuspended in 1 mM Tris-EDTA solution. We analysed data from 8 combinations with at least 2 technical replicates for each infection combination and 2 controls with 3 technical replicates.

After successfully extracting and purifying RNA from both the host and the bacterial samples involved in the infection, 48 samples underwent ribodepletion using the RiboZero Gold kit (Illumina, San Diego, USA). The library preparation was elaborated using the NEBNext® SingleCell/Low Input cDNA Synthesis kit (New England Biolabs). The prepared libraries were then sequenced, utilising an S4 150 bp configuration on the Illumina NovaSeq 6000 platform (for sequenced samples see Table S3). The obtained reads are available under the bioproject IDs PRJNA1124259 and PRJNA1124539.

### RNA-seq analysis

Read quality was evaluated with the BBDuk function from BBMap v. 39.06 [30], setting a minimum length at 35 bp and a minimum quality of 35. This step ensured the preservation of the longest, high-quality reads while discarding low-quality segments. The read quality was also evaluated before and after trimming using FastQC v. 0.12.1 [31].

We aligned the filtered reads to their respective reference genomes using STAR v. 2.7.11b [32]. Reads from infected THP-1 macrophages were mapped to *Homo sapiens* (v 104, accession number GCA_000001405.15), while those from infected BoMac macrophages were aligned to *Bos taurus* genome (v 109.12, accession number GCA_002263795.2).

The following analysis was conducted using R v. 4.3.2, with the code available in <u>PathoGenOmics-Lab/CrossInfectMTB</u>. Gene expression quantification was performed using featureCounts function from R package Rsubread v. 2.16.1 [33]. This step involved mapping the high-quality reads to genomic features from the Ensembl annotation (109 releases for *B. taurus* and 104 for *H. sapiens*). We filtered the raw gene expression counts using the HTSfilter R package v. 1.42.0 [34] to eliminate genes with non-informative signals. Normalisation of the raw counts was then carried out using the DESeq2 R package v. 1.42.0 [35], based on the mean ratios method. Once we obtained the sequencing we evaluated the expression of the genes, and one sample that clustered with controls was removed.

### Differential expression analysis

Differentially expressed genes (DEG) between the conditions of interest (comparisons detailed in Figure S1D) were obtained with DESeq2 analysis. To identify common DEGs between *H. sapiens* and *B. taurus*, we employed the biomaRt R package v. 2.58.0 [36], and we kept one-to-one orthologs. Each contrast’s significantly differentially expressed genes were determined using thresholds of a p-value < 0.05 and a log-fold change > 1 or < - 1. Gene Ontology Enrichment Analysis (GOEA) and KEGG Pathways Enrichment Analysis (KEA) were performed to identify the most enriched Gene Ontologies (GOs) and KEGG terms. The enrichment analysis of DEGs was analysed with the clusterProfiler R package v. 4.10.0 [37] and the AnnotationDbi R package v. 1.64.1, for gene enrichment we employed the org.Hs.eg.db and org.Bt.eg.db v. 3.19.1 R packages. The results from GOEA were streamlined and visualised using the rrvgo R package v. 1.14.1 [38]. Additionally, all analyses across our study were visualised comprehensively using the ggplot2 R package v. 3.5.0.

Correlation of differentially expressed genes and infection ratios were determined by a custom R script applying a linear model with R core basic functions, results were visualised with ggplot R package v. 3.5.0.

### Differential exon usage analysis

Once the reads were aligned we quantified the reads using the python library HTSeq [39] v.

2.0.5. Then, the differential exon usage (DEU) was assessed with the DexSeq v. 1.50.0 [40] R package. The steps involving alternative splicing analysis were executed with the pipeline of nf-core/rnasplice v. 1.0.3. Once the differential exon usage was assessed, the contrasts between the comparisons of interest were analysed with a custom R package and the DEUs were visualised with ggplot2 v. 3.5.0 and DexSeq v. 1.50.0 [40] packages. The genes containing the DEUs were analysed with a gene enrichment analysis for GOEA and KEA by the clusterProfiler R package v. 4.10.0 [37] and the AnnotationDbi R package v. 1.64.1 functions.

### RT-qPCR assays of *plcABCD* genes

We quantified the expression of the four phospholipase C genes (*plcA*, *plcB*, *plcC,* and *plcD* respectively, see Table S4) by qPCR relative to *sigA*. For primer details, see Table S5.

In the retro transcription analysis, approximately 10 μL of RNA from *M. tuberculosis* derived from in vitro infections was denatured at 70°C for 10 minutes. The RT reaction mix, consisting of 6 μL of 5x First Strand PCR buffer, 4 μL of 25 mM dNTPs, 6 μL of 25 mM MgCl2, 1 μL of 1 mg/mL random primers, 0.5 μL SuperScript II reverse transcriptase, and 22.5 μL of RNase-free water, was then added to the RNA. This mixture was incubated at 25°C for 10 minutes, followed by 1 hour at 42°C for the reverse transcription.

For the qPCR reactions, the following components were used: 10 μL SYBR Green, 2 μL of 10 μM primer mix, 6 μL of MilliQ H_2_O, and 2 μL of template DNA/cDNA. The qPCR analysis was performed using a QuantStudio™ 3 Real-Time PCR System (ThermoFisher Scientific) with the following protocol: 1) Initial denaturation at 50°C for 2 minutes and 95°C for 10 minutes. 2) 40 cycles of annealing/extension at 95°C for 15 seconds and with a ramp from 95°C to 65°C at 1.6°C/s, followed by 65°C for 1 minute. 3) Final disassociation at 95°C for 15 seconds.

### *plcABCD* expression analysis

To assess the expression of the phospholipase C genes using qPCR, genomic DNA from a chimpanzee bacillus strain was used as a positive control at a concentration of 3·10^6^ genomes per qPCR reaction. The genome concentration was extrapolated based on the genome size of the H37Rv strain, following quantification of the genomic DNA extract using Qubit. For each case (combination of bacterial strain and macrophage type), three replicates were selected, focusing on those with the lowest Ct value for the *sigA* gene.

To calculate the relative expression of each gene, standard curves were generated using the genomic DNA of the chimpanzee bacillus as a template, since it contains the four phospholipase C genes. The number of copies in the qPCR reaction was inferred from the Qubit quantification and genome size. A logarithmic series of descending concentrations, starting from 3·10^6^ down to 0 copies, was created for each gene in each qPCR reaction.

A linear model was then fitted to the log10(template concentration), and the amplification efficiency (E) for each primer was determined by:

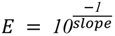

To determine the relative expression of each gene to *sigA*, we used the efficiencies of each primer and the Ct values, calculated by:

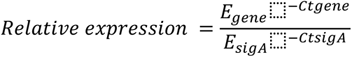

We normalised the variation of the relative expression given by replicates, where CtsigA_C_ is the Ct value of *sigA* amplification reaction of the positive control, with:

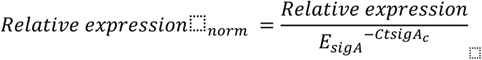

## Results

### *In vitro* infection ratio matches with preferences by host

To assess the compatibility between *M. tuberculosis* lineages with their host cells, we compared infection ratios across the four bacterial-host combinations at 4 hours post-infection (hpi), 1 day, and 3 days post-infection (dpi). We could detect that L5 showed a significantly larger median infection ratio in human THP-1 cells than in BoMac (48.33% vs. 12.85%; Mann-Whitney U test p-value = 0.00017, Figure 1A), while *M. bovis* showed significantly larger median infection ratios in bovine BoMac cells than in THP-1 (47.05% vs. 22.87%, Mann-Whitney U test p-value = 0.043, Figure 1A) at 3 dpi. Interestingly, L6 and the Chimpanzee bacillus did not show significant differences in infection ratio between THP-1 and BoMac (L6: 34.68 vs. 18.75 respectively; *M. bovis*: 28.07 vs. 20.05 respectively, Figure 1A).

**Figure 1.**
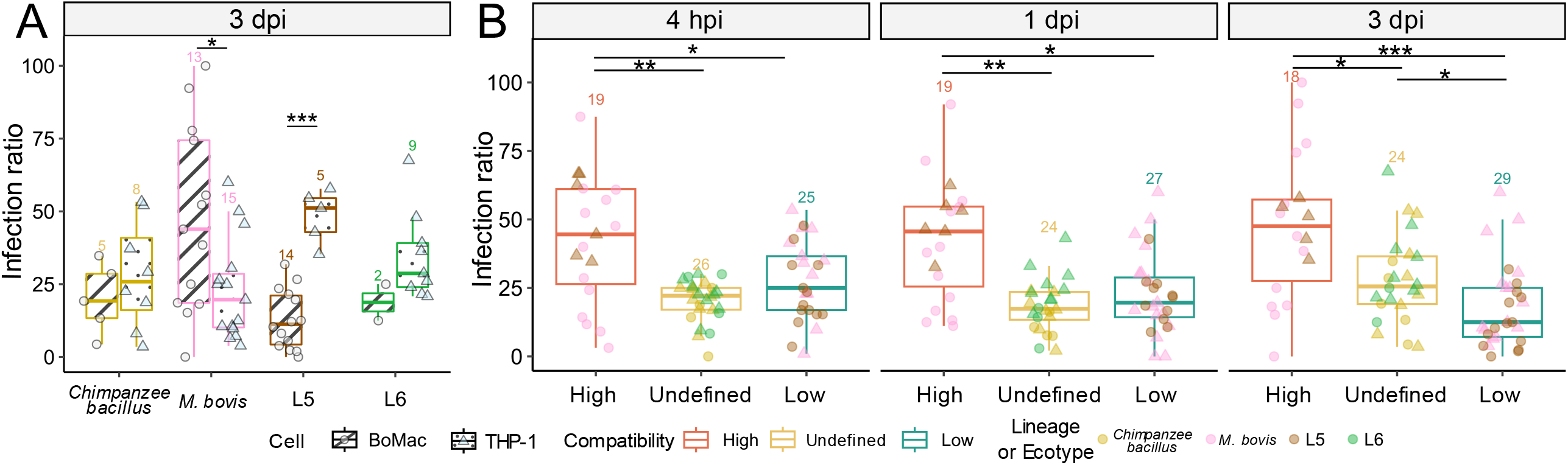
Infectivity variability assessed by microscopy infection ratio in cross-infections of THP-1 and BoMac cells. **A)** Differences in infection ratios between cells infected with the same lineage at 3 dpi, evaluated by the Mann-Whitney U test. **B)** Differences in infection ratios for each compatibility group at different times post-infection were evaluated using the Dunn test with the Benjamini-Hochberg correction. In both panels, the number of replicates per condition is indicated above each group and significant differences are shown as *: p ≤ 0.05; **: p ≤ 0.01; ***: p ≤ 0.001; ****: p ≤ 0.0001.

When we group the high compatibility combinations (*M. bovis* in BoMac and L5 in THP-1) and compare it with the lower compatibility (*M. bovis* in THP-1 and L5 in BoMac), or with the infections of the Chimpanzee bacillus and L6 for which the compatibility is not as clear (undefined from now on), we detected that high compatibility infections showed significantly larger infection ratios than the rest (Figure 1B) at all time points. Specifically, at 3 dpi, the undefined compatibility infections showed an intermediate infection ratio, being significantly smaller than the high compatibility combinations (28.10% vs. 47.40%, Dunn test padj-value = 0.045; Figure 1B) and significantly larger than the low compatibility infections (28.10% vs. 18.03%, Dunn test padj-value = 0.045; Figure 1B).

We also explored the effects of infection on cell death by flow cytometry analysis categorising outcomes into necrosis, apoptosis, and live cells. THP-1 displayed larger apoptosis ratios than BoMac cells independently of the infecting strain (Figure S2A). We could not detect significant differences in cell death between high and low-compatibility infections, but both showed significantly smaller proportions of necrosis than the undefined compatibility infections (Figure S2B). However, although the magnitude of the difference is small and not significant, we could detect a tendency where high compatibility infections tend to show a slightly larger necrosis proportion than the low compatibility (2.52 vs. 1.50 Figure S2B). This can be also observed when comparing necrosis between both cell types (Figure S2A), where *M. bovis* showed higher necrosis in BoMac cells than in THP-1 (3.04 vs. 1.43) and L5 in THP-1 vs. BoMac (2.11 vs. 1.56).

### Transcriptional signatures of low and high compatibility infections

Global transcription profiles show different host responses between high and low compatibility infections, especially in THP-1 where the overall variance structure of the global transcriptome separates better the infections with L5 and *M. bovis* (Figure 2A and 2B). To explore expression differences due to host compatibility, we performed a differential expression analysis of orthologous genes between high and low compatibilities (Figure 2C). When comparing orthologous genes between both cell types, high compatibility infections were characterised by a common enrichment of transcription in genes related to cell proliferation, including cell cycle regulation and DNA replication activities within the cell nucleolus in both cells (137 differential expressed orthologs genes (DEOGs) upregulated, log_2_ fold change > 0 and p-adj < 0.05; Table S6, Figure 2C, 2D and 2E). Biological processes identified by gene ontologies (GOs) included functions involved in cell cycle regulation and DNA replication activities within the cell nucleolus (Figure 3A and 3B, Table S7). Molecular functions included damaged DNA binding, catalytic activity acting on DNA, and helicase activity (Table S7). Enriched cellular components included the condensed chromosome, midbody, nuclear envelope, centriole, and oxidoreductase complex (Table S7). KEGG database enrichment analysis revealed an enrichment of pathways related to the cell cycle and Fanconi anemia pathways (Table S7).

**Figure 2.**
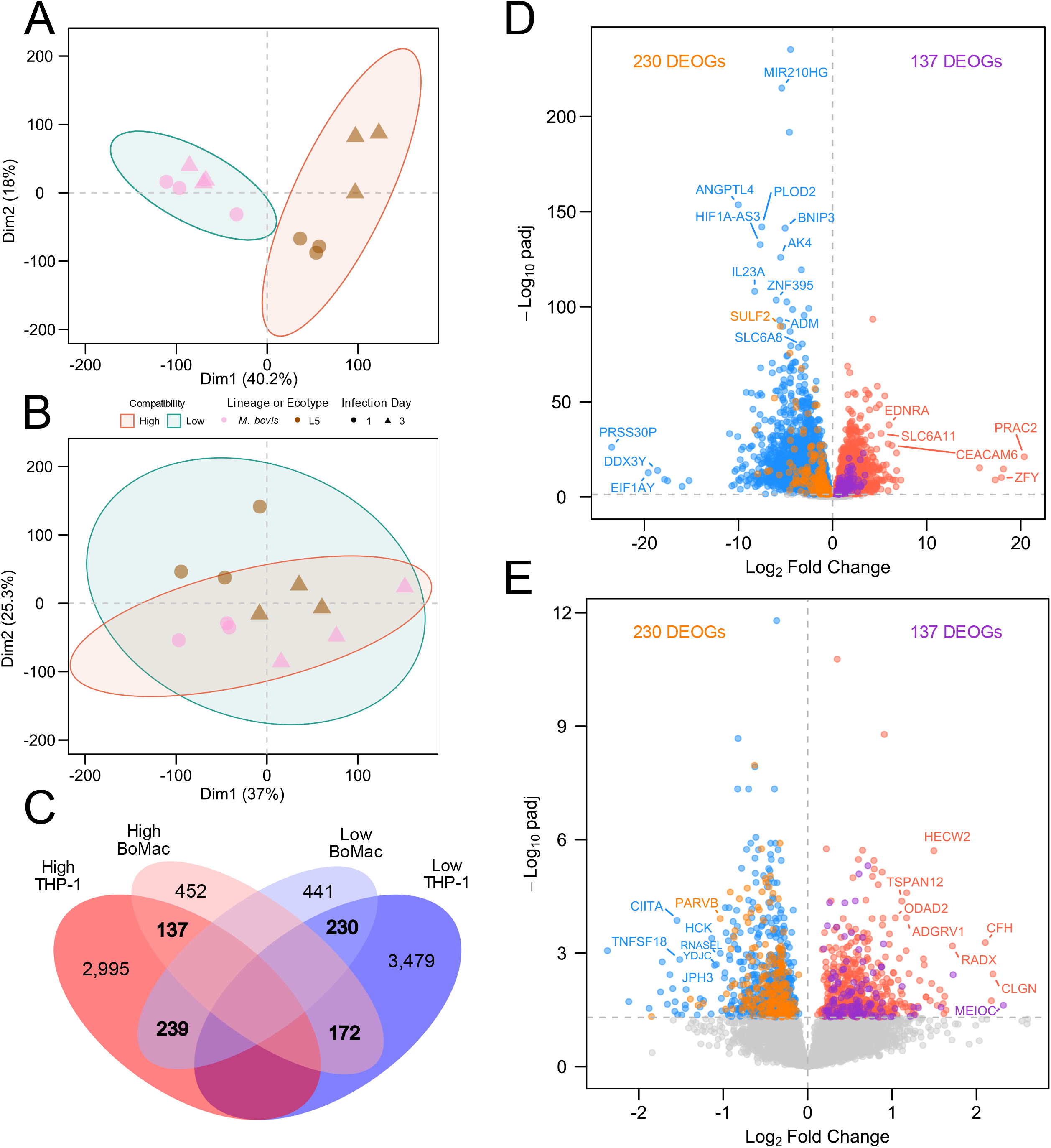
Differential host gene expression contrasting high vs. low compatibilities. Dimensionality of gene expression for all strains infecting THP-1 cells **(A)** or BoMac cells **(B)**. **C)** Common and unique DEOGs for THP-1 and BoMac cells in the contrast of high vs. low compatibility. Volcano plot of differentially expressed genes in the contrast of high vs. low compatibility in THP-1 cells **(D)** and BoMac cells **(E)**. Gene labels are shown only when p-adj < 0.05 and log_2_ fold change > 5 or < −5 for THP-1 cells and log_2_ fold change > 1 or < −1 for BoMac.

**Figure 3.**
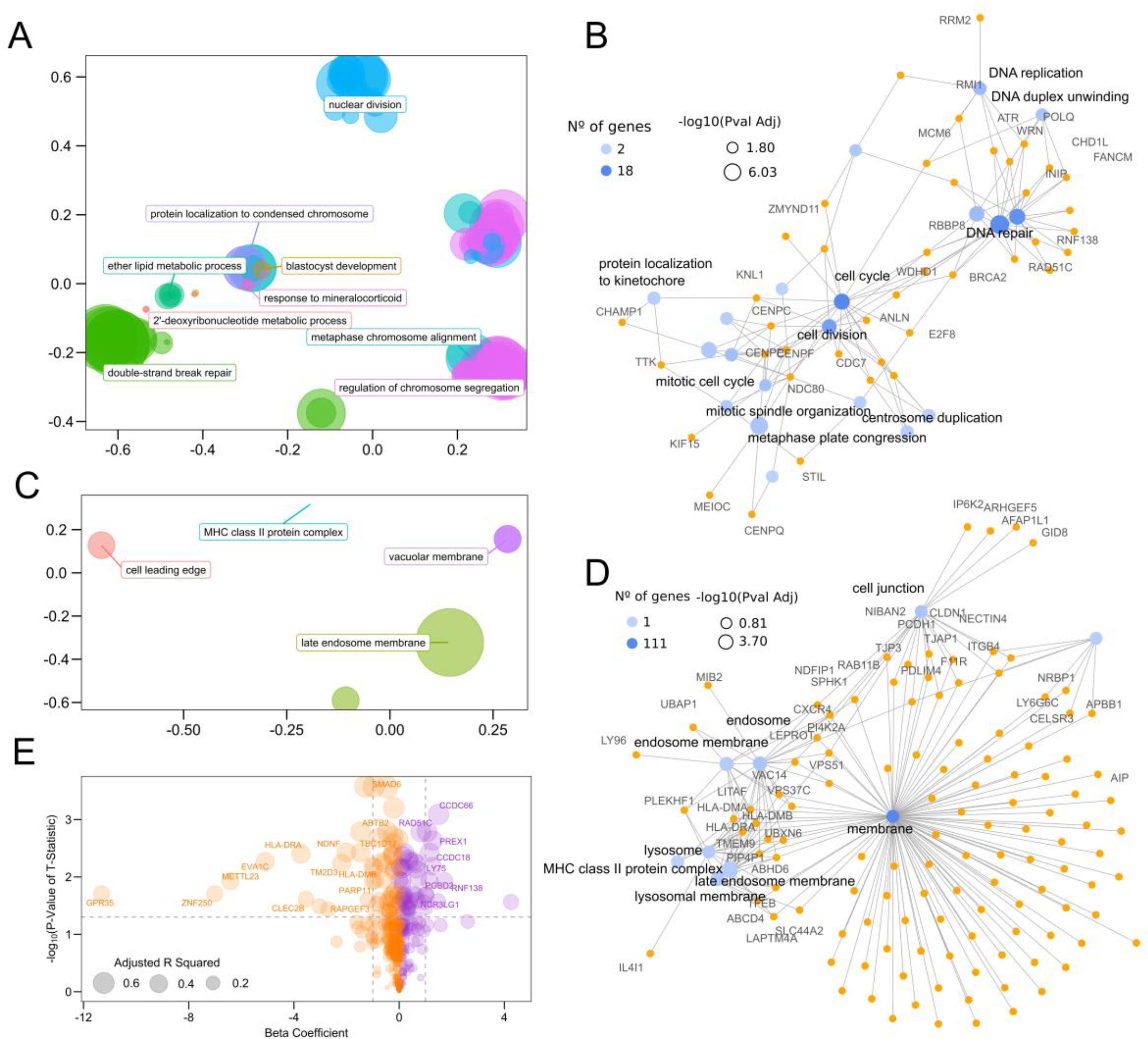
Gene expression enrichment of the host in the contrast of high vs. low compatibilities. **A)** Biological processes enriched among the 137 DEOGs upregulated in high vs. low compatibility **B)** Top 20 biological processes enriched in 137 DEOGs upregulated in high vs. low compatibility for high compatibility **C)** Cellular components ontology of 230 DEOGs downregulated in high vs. low compatibility. **D)** Top 10 cellular components ontology of 230 DEOGs down regulated in high vs. low compatibility. **E)** Correlation between infection ratio and expression of the 137 upregulated DEOGs (purple) and 230 downregulated DEOGs (orange). The distances between points in panels A and C represent the similarity between terms, and the size of the point represents the scores of the GO term.

But when we stratified high compatibility infections by cell type, we could detect certain genes exclusively found in one host but not the other. High compatibility THP-1 infection was characterised by increased transcription of 2,456 differential expressed genes (DEG) (log_2_ fold change > 1 and p-adj < 0.05, Table S8) that are enriched in KEGG pathways related to replication and repair: the cell cycle, DNA replication, cell growth and death, nucleotide, amino acid, and carbohydrate metabolism (Table S9). Upregulated biological processes included mainly functions related to chromosome segregation, DNA replication, cell division, and responses to DNA damage. High compatibility BoMac infections also include upregulated functions related to cell cycle, DNA replication, helicase, and ATP activity (758 DEGs log_2_ fold change > 0 and p-adj < 0.05, Table S8).

Low compatibility infections were characterised by an enrichment of transcription in genes related to intracellular bacterial containment and immune evasion (230 DEOGs log_2_ fold change < 0 and p-adj < 0.05; Table S6, Figure 2C, 2D and 2E). The enrichment of cellular components included the late endosome membrane, vacuolar membrane, MHC class II protein complex, and cell leading edge, (Figure 3C and 3D; Table S7). Again, if we stratify by cell line, low compatibility THP-1 infection was characterised by an upregulation of 3,654 DEGs (log_2_ fold change < −1 and p-adj < 0.05; Table S8). Those genes were enriched in processes of immune systems such as signalling molecules, cellular interaction and transduction, regulation of T cell activation, cytokines, and interleukins among others. Additionally, lipid metabolism and biological functions like oxygen regulation within the cell were also observed (Table S9). However, high compatibility BoMac infections showed 910 DEGs (log_2_ fold change < 0 and p-adj < 0.05; Table S8) related to antigen processing and presentation, and T helper differentiation (Table S9).

### Strain-specific transcriptional signatures

When we compare expression profiles between strains independently of the host cell, we observed 239 DEOGs commonly upregulated in L5 strain infections (Figure 2C, Table S6). L5-specific transcription compared to *M. bovis* was enriched in genes related to peroxisome, microbody, vesicle tethering complex, and chromosomal and telomeric regions (Table S7). However, *M. bovis*-specific upregulated genes compared to L5 (170 DEOGs; Figure 2C, Table S6) were enriched for the KEGGs pathways related to the ferroptosis pathway, and molecular functions related to ubiquitin-like protein ligase activity, cadherin binding and rough endoplasmic reticulum membrane (Table S7).

### Temporal transcriptional signatures depend on compatibility

We also observed transcriptional differences at different stages of infection between high and low compatibility. At early stages, high compatibility infections were characterised by increased transcription in KEGG pathways that include signal transduction, infectious disease, focal adhesion, and metabolism of terpenoids and polyketides (528 DEOGs, Table S10, Table S11). Enriched biological processes involved ribosome biogenesis, RNA processing, and cellular regulations such as negative regulation of apoptotic signalling pathways (Table S11). Conversely, low compatibility infections were characterised by enriched KEGG pathways in pentose phosphate, thermogenesis, and oxidative phosphorylation among others (367 DEOGs, Table S10, Table S11). In the biological process, we observe the ribose phosphate biosynthetic process, energy derivation by oxidation of organic compounds, regulation of cellular component size, response to oxygen levels and oxidative stress, regulation of leukocyte apoptotic process, and vesicle-mediated transport between endosomal compartments (Table S11).

At later stages of infection (3 dpi), high compatibility (638 DEOGs, Table S10) was characterised by enriched transcription of KEGG pathways related to lysosome, phosphatidylinositol signalling, inositol phosphate metabolism, DNA replication, and glycan degradation (Table S11). In the biological processes, we obtained enriched functions related to liposaccharide metabolic processes, chromosome segregation, and cellular reorganisation such as nuclear division, and autophagosome organisation. Later stages in low compatibility infections (266 DEOGs, Table S10) were characterised by an enriched transcription in molecular functions enriched for protein lysine deacetylase activity (Table S11).

### Molecular signatures of infection ratio by host compatibility

To decipher which genes could explain the lower *in vitro* infection ratio of the high compatibility infections, we correlated the expression levels of selected genes with infection ratios (Table S12). We found 10 genes significantly negatively correlated with lower infection ratio, which include 3 genes related to MHC II antigen presentation: *CLEC2B*, *HLA-DRA*, and *HLA-DMB* (Figure 3E). Conversely, we found 48 genes significantly positively correlated to a high infection ratio, which include genes involved in double-strand break repair via homologous recombination and meiotic nuclear division (Figure 3E). Among the strongest correlations with high infection ratios we found *NCR3LG1* which is a Natural Killer Cell Cytotoxicity Receptor 3 Ligand, and the gene *RNF138* which is related to MHC I and innate immune system.

### Differential exon usage by host compatibility

In both, high and low compatibilities, we have identified different uses of exons leading to different isoforms in THP-1. We observed 982 genes (p-adj < 0.05, log_2_ fold change > 1 and < −1) that harboured some exons more expressed in high-compatible (L5 strain) infections and other exons in low-compatible (*M. bovis* strain) infections (Figure 4A, Table S13). These genes include enriched functions of cell growth, organisation, and mobility, but also GTPase activity (Figure 4B-D). In the KEGG database, we observed enriched pathways in the ECM-receptor interaction, focal adhesion, and cytoskeleton in muscle cells (Table S14). *IL-16* shows 28 exons with 15 differential exon usages (DEUs) (p-adj < 0.05), exon E025 and E019 appeared upregulated in *M. bovis*, and 13 other exons were upregulated in L5 (Figure 4I; Table S13). Another gene that we observed is *SIGIRR* related to the negative regulator of the Toll-like and IL-1R receptor signalling pathways. We observed 17 exons in the gene with 14 DEUs, 11 of them upregulated in L5 and 3 upregulated in *M. bovis* (Figure 4J, Table S13). Gene *PRKCD* encodes a protein that can positively or negatively regulate apoptosis, and we found 23 exons with 6 DEUs where 3 exons were upregulated in L5 infections and another 3 upregulated in *M. bovis* (Figure 4K; Table S13).

**Figure 4.**
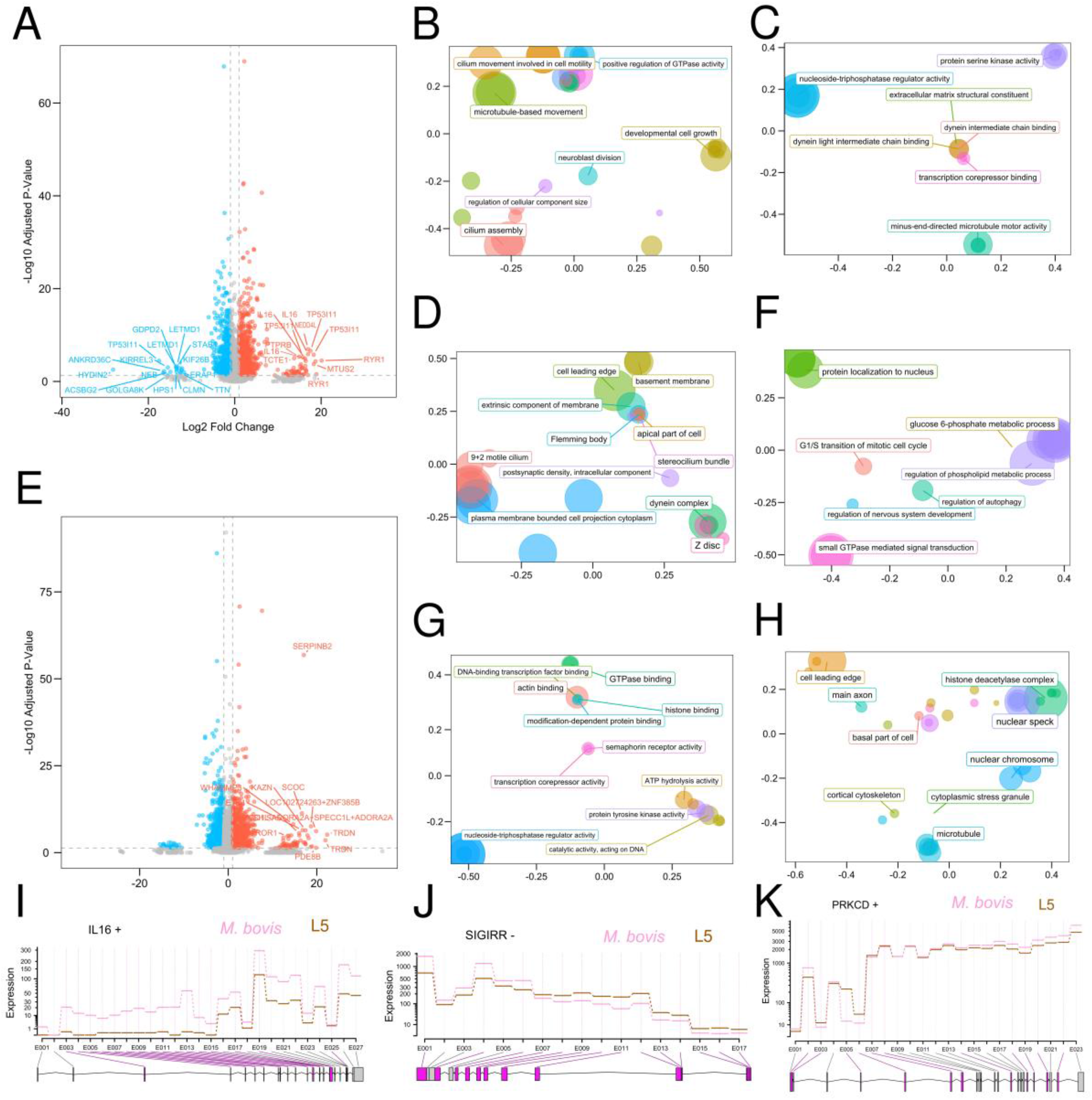
Differential exon usage for the high and low compatibility infections. **A)** Differential exon usage for genes where blue dots are exons expressed in high compatibility infections and red dots are other exons of the same gene expressed in low compatibility infections. Functional enrichment for genes in panel A for biological processes **(B)** molecular functions **(C)** and cellular components **(D). E)** Differential exon usage for genes where blue dots represent exons of genes only expressed in high compatibility infections and red dots exons of genes only expressed in low compatibility infections. Functional enrichment for genes related to high compatibility in panel E for biological processes **(F)** molecular functions **(G)** and cellular components **(H)**. The distances between points in panels B-C and F-H represent the similarity between terms, and the size of the point represents the scores of the GO term. Fitted expression values of each of the exons in the genes *IL-16* **(I)**, *SIGIRR* **(J)** and *PRKCD* **(K).** For panels I, J, and K, exons with a differential usage between high (L5 strain) and low (*M. bovis* strain) are highlighted with purple.

Additionally, we detected 2,379 genes (p-adj < 0.05, log_2_ fold change < −1, Figure 4B, Table S13) with all their exons more expressed in L5 infections (Table S13). They include biological processes related to the regulation of autophagy and phospholipid metabolic process, protein localization to the nucleus, G1/S transition of mitotic cell cycle, GTPase mediated signal transduction and glucose 6-phosphate metabolic process (Figure 4F). For the molecular functions, we observed actin and DNA binding (Figure 4G), and in the cellular components, we observed functions related to the organisation of the cell (Figure 4H; Table S14).

Finally, we found 1,749 genes (p-adj < 0.05, log_2_ fold change > 1, Figure 4B, Table S13) where all their exons were more expressed in *M. bovis* infection (Table S13). In this category, we observed biological processes such as regulation of endocytosis, peptidyl-serine phosphorylation and modification, and molecular functions like GTPase regulator activity, nucleoside-triphosphatase regulator activity, and ATP hydrolysis activity. Cellular components were related to cleavage furrow, histone acetyltransferase complex, mitotic complex, and nucleotide-excision repair complex. In the KEGG database, we observed pathways related to DNA replication, Fc gamma R-mediated phagocytosis, biosynthesis of cofactors, and VEGF signalling pathway, among others (Table S14).

### Correlation between the signature virulence genes of *M. tuberculosis* and host receptors

Due to the potential role of *plc* in the virulence of *M. tuberculosis* and its differential context between human and associated lineages, we studied the expression of these genes, and its correlation with the *PTGER1*, *PTGER2*, *PTGER3*, and *PTGER4* receptors. When comparing the expression of different strains, we found that the L6 strain showed almost negligible expression of *plcC* during infection independently of the infected cell (36.05 L6 vs. 2084.10 L5, Dunn test adjusted p-value = 0.001; 36.05 L6 vs. 1368.92 *Chimpanzee bacillus*, adjusted p-value = 0.039; Figure S3A). Additionally, when comparing the expression of the same strain between cell lines, we found that *plcD* expression was significantly smaller in L5 infections in high compatibility cells (THP-1) compared to low compatibility cells (BoMac) (2838.35 THP-1 vs. 4676.58 BoMac, t-test p-value = 0.00121, see Figure S3B).

*PTGER* receptors exhibited significantly higher expression when infected with the least compatible strains such as THP-1 infected with *M. bovis* strain (all *PTGER*; Figure 5A) and BoMac with L5 strain (*PTGER3*; Figure 5A). *PTGER3* was also more expressed in THP-1 cells than in BoMac cells when infected with L6, but not with the chimpanzee bacillus. Additionally, *PTGER2* showed significantly increased expression in THP-1 than in BoMac at 3 dpi when infected with the undefined compatibility strains L6 and the chimpanzee bacillus (Figure 5A).

**Figure 5.**
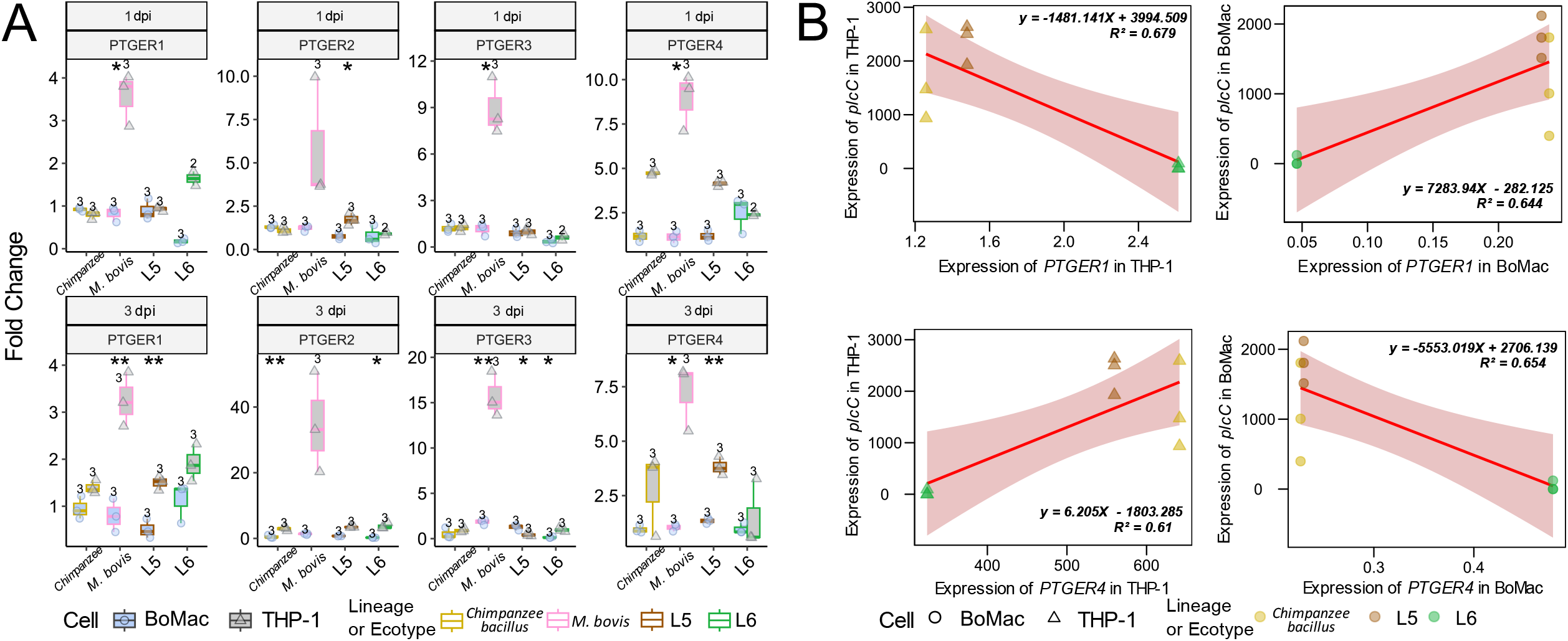
Expression of *M. tuberculosis* phospholipases C and *PTGER* genes in BoMac and THP-1 cells. **A)** Comparison of *PTGER* genes expression between BoMac and THP-1 cells, evaluated by the T-test. Significant results are shown as *: p ≤ 0.05; **: p ≤ 0.01; ***: p ≤ 0.001; ****: p ≤ 0.0001. Fold change was calculated by comparing sample values to the control mean. **B)** Correlation between bacterial (*plcC*) and host (*PTGER1* and *PTGER4*) gene expression in BoMac and THP-1 infections at 1 dpi.

To explore the interactions between host and bacterial expression, we performed correlations between the expression of *plc’s* and *PTGER* genes. A significant negative correlation was observed between all *PTGER* receptors and *plcA* and *plcB* expression, and also between *PTGER2* and *PTGER3* with *plcC* expression (Figure S3C). This correlation is mainly driven by the absence of the three phospholipases in *M. bovis,* which led to no expression of the gene and correlated with a very high expression of the *PTGER* genes. To understand the correlation beyond the effect of *M. bovis*, we examined the correlation patterns between THP-1 and BoMac cells excluding *M. bovis* infections. Significative positive correlation was detected in *plcC* and *PTGER1* expression (p-value = 0.00630) and negative with *PTGER4* expression (p-value = 0.0129; Figure 5B) in THP-1 cells. The opposite pattern was observed in BoMac cells (p-value = 0.00925 for *PTGER1*; p-value = 0.00832 for *PTGER4*; Figure 5B). This correlation seems strongly driven by the low expression of plc genes in the L6 strain described above.

## Discussion

Pathogens have evolved to interact with specific molecules in the host’s cells, like cell receptors, nutrients, toxics, or specific immune molecules that determine the molecular determinants of host-pathogen compatibility. In the case of *M. tuberculosis*, one central aspect is why specific ecotypes or lineages affect particular hosts, and if there are molecular determinants of such specificity. There is a wealth of data providing host transcriptional responses to different *M. tuberculosis* strains during *in vitro* infection [41,42]. However, our approach focuses on comparing specific patterns of different predicted compatibility between strain and host, allowing us to decipher common molecular signals of different host cells to the most compatible infection and specific patterns of certain strains independently of the host.

We confirmed our hypothesis that bacteria infecting the most compatible host cell display a higher infection ratio associated with more infectivity in an *in vitro* system. But bacteria like the Chimpanzee bacillus, which has been isolated only once, and their host preference is not known, showed intermediate patterns of infection ratio between the other two strains and similar infection ratio between both hosts. Similar patterns were detected in the L6 strain, despite belonging to a human-associated lineage. L6 is predominantly isolated from humans, but their association with immunocompromised individuals, their phylogenetic position among animal-associated lineages, and its larger genomic diversity led to the hypothesis that the L6 strain will show a different pattern compared to L5 a human-associated strain [11,22]. Interestingly, L6 is the closest lineage to lineage A1, which includes the Chimpanzee bacillus [11,22]. Our results confirm that the L6 strain and the Chimpanzee bacillus behave more similarly in *in vitro* infections than with the other strain from a human-associated lineage (L5).

In order to decipher the molecular determinants of *in vitro* compatibility of different strains in human and bovine cells, we compared the expression profiles. Comparative genomic studies between *Bos taurus* and *Homo sapiens* face significant challenges due to differences in genome reference quality and completeness. *B. taurus* genomes, despite improvements, still have inconsistencies and gaps [43], and these disparities between references complicate RNA-seq comparative analysis between species because the comparison of DEGs, and even more splicing events are challenging. Despite this, we were able to focus on retrieving common transcriptional patterns of high and low compatibility *in vitro* infections. We found that bacteria infecting the most compatible host cell could potentially evade the immune system’s weaponry because it inhibits genes related to antigen presentation (MHC II), cellular compartmentalization, and endosome creation, which are under-expressed. These pathways might play a crucial role in *M. tuberculosis* infections, as the bacteria’s ability to inhibit antigen presentation and manipulate host cellular processes allows it to evade immune detection and establish chronic infections [44,45]. On the contrary, the most compatible infections result in upregulation of the host’s cell replication and DNA, which might be interpreted as a bacterial strategy to increase cell replication and thus spread the infection. This strategy was shown for *Chlamydia trachomatis* in an *in vitro* infection system that causes the proliferation of host cells to maintain a persistent infection [46]. Conversely, strains infecting their least compatible host cell display the reverse pattern. The host overexpressed compartmentalization, and specific genes such as *UBXN6*, *ABCD4,* or *TFEB* among others could lead to the activation and elimination of infected cells. The importance of these pathways in *M. tuberculosi*s infections has been highlighted in studies showing that effective compartmentalization in phagosomes and endosomes, and immune activation can restrict bacterial growth and promote clearance of the infection [47]. In addition to common signatures, we could detect strain-specific patterns such as the upregulation of ferroptosis pathways in *M. bovis* infections compared to the infection with the L5 strain. Ferroptosis, characterised by iron-dependent lipid peroxidation, has been shown to be a critical mechanism of necrosis during *M. tuberculosis* infection, both *in vitro* and *in vivo* [48].

Additionally, we found major differences in the temporal response of the most and least compatible infections. Infection with the most compatible strains, results in a nonspecific early transcriptional response towards infection, but is directed to cellular reorganisation by upregulating genes involved in cellular metabolism and ribosome generation, impairing apoptosis signalling and promoting cellular mobilisation in terms of cell adhesion. At later stages, these strains upregulate genes related to pathways to manipulate lysosomes and organise autophagosomes for survival [49], which are important in the control of *M. tuberculosis* infection [50]. Additionally, the transcriptional profile suggests these strains might specialise in resource acquisition by degrading glycans and activating polysaccharide metabolism. This is consistent with a previous study that demonstrated *M. tuberculosis* accesses glucose *in vivo* and relies on glucose phosphorylation for survival during chronic mouse infections [51]. In contrast, infections with the least compatible strains result in more specific anti-pathogen transcriptional responses early on, such as upregulate genes that modify oxygen levels and oxidative stress, potentially contributing to the antimycobacterial function of peroxisomes in the cytosol of human macrophages [52], as well as activating vesicle transport in endosomal compartments to control the bacteria [53].

But other transcriptional signatures, such as alternative splicing can also play an important role in virulence-specific responses, as previously found during *in vitro* infections with virulent and avirulent laboratory *M. tuberculosis* strains [54]. In our study, we observed hundreds of genes with different isoforms associated with low or high-compatibility THP-1 infections, and although we cannot precisely determine how these isoforms affect infection, we can identify patterns associated with compatibility that could be investigated in the future. We detected different isoforms of *IL-16* in high vs. low compatibility THP-1 infections, where several exon usages were related to low compatibility infections. *IL-16* functions as a chemoattractant modulating T cell activation [55] and the different isoforms observed could potentially affect gene functionalities and impair the immune response.

Genomic differences between human-associated lineages and animal-associated ecotypes could reveal the nature of the molecular differences implicated in host specificity, and one of the clearest distinctions in gene content between human-associated and animal-associated ecotypes is RD5 [23,56]. RD5 has been recurrently lost in animal lineages, and it contains three out of four *plc* genes and *PPE* genes [56]. We found significant differences in the expression of *plc* genes between different strain-host combinations. For instance, the negligible expression of *plcC* in L6 infections at 1 dpi highlights the unique expression dynamics of this strain, which presents a non-synonymous mutation in the *plcC* gene (accession number: ERR2704688). Previous studies have demonstrated that the *plc*’s inhibits the expression of *PTGER2* and *PTGER4* [26] which affects immune response and inflammation through the eicosanoid pathway [57]. We have confirmed the association of plc’s and *PTGER* expression. But we also found higher expression of *PTGER* when infected with the least compatible strains, which might indicate a response towards controlling the infection more efficiently. Additionally, the opposite patterns in the correlation of expression of *plcC* and *PTGER* between THP-1 and BoMac cells could suggest that host cell type significantly influences the interaction between *M. tuberculosis* virulence factors and host receptors, potentially impacting infection outcomes.

Our results are limited by our experimental setup. We use human-associated L5 and L6 strains and animal-associated chimpanzee bacillus and *M. bovis* strains, which could not be representative of broader patterns across the whole *M. tuberculosis* lineage. Variability can be expected among different strains within the same lineage and among lineages [58,59]. Therefore, additional studies with more strains of the same lineage are needed to generalise our findings. Another limitation of our study is the use of THP-1 and BoMac cell lines, which may exhibit altered gene expression profiles, signalling pathways, and functional characteristics compared to primary cells [60]. Therefore, the investigation of the cellular and molecular responses of primary immune cells could provide a more realistic picture of the complexity of the immune system, including differences between host populations.

Overall, our findings have confirmed our hypothesis that more compatible strain-host combinations result in a higher infection ratio and non-efficient molecular response of the pathogen from early on the infection. This contributes to a deeper understanding of the molecular response of human and bovine cell lines challenged with different *M. tuberculosis* strains in the context of the complex interplay between strain-host compatibility.

## Acknowledgments

We extend our gratitude to Prof. Sébastien Gagneux and Dr. Iñaki Comas for generously providing the mycobacterial strains, Dr. Raquel Conde and Dr. Esther Julián for their provision of THP-1 cells, and Dr. Matthias Schweizer for sharing the BoMac cells.

The computational analyses were performed on the HPC cluster Garnatxa at Institute for Integrative Systems Biology (I^2^SysBio).

## Data availability

The code for the bioinformatic methods and analyses described are available at https://github.com/PathoGenOmics-Lab/CrossInfectMTB.

## Funding

This work was financially supported by project PID2021-123443OB-I00 funded by MCIN/AEI/10.13039/501100011033 and by FEDER A way to make Europe. PRR is supported by CNS2022-135116 funded by MCIN/AEI /10.13039/501100011033 and by the European Union NextGenerationEU/PRTR. MCG is supported by a predoctoral contract from the Generalitat Valenciana (CIACIF/2021/159) and Project CIPROM2021-053 from Generalitat Valenciana.

## Disclosure statement

The authors declare no conflict of interest.

## Author Contributions

**PRR:** data curation, formal analysis, investigation, methodology, software, validation, visualisation, writing - original draft. **MCG:** investigation, validation, methodology, writing - original draft. **GS:** formal analysis, investigation, methodology, writing – review & editing. **HH:** investigation, writing – review & editing. **MC:** conceptualization, funding acquisition, investigation, methodology, project administration, supervision, writing - original draft.

---

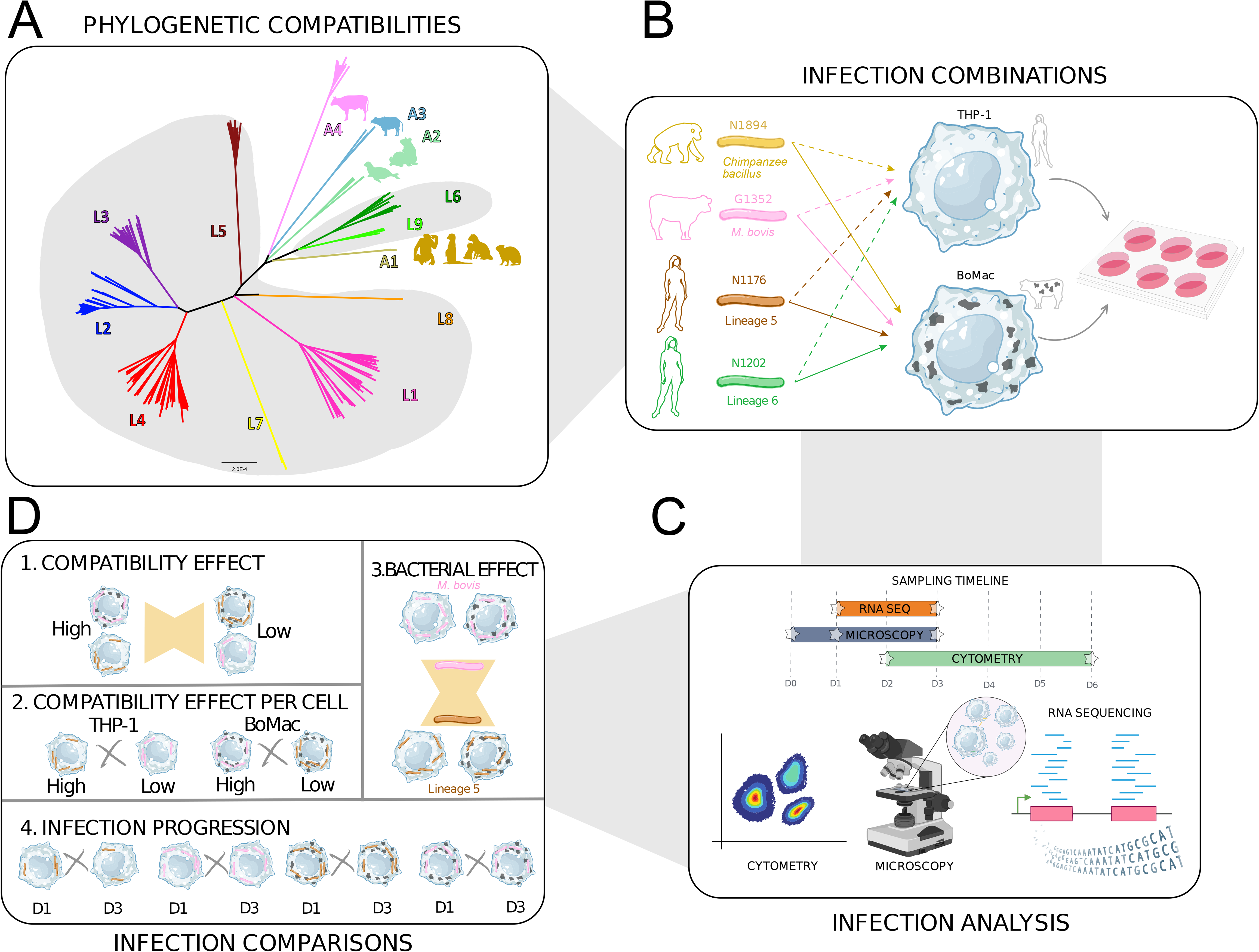
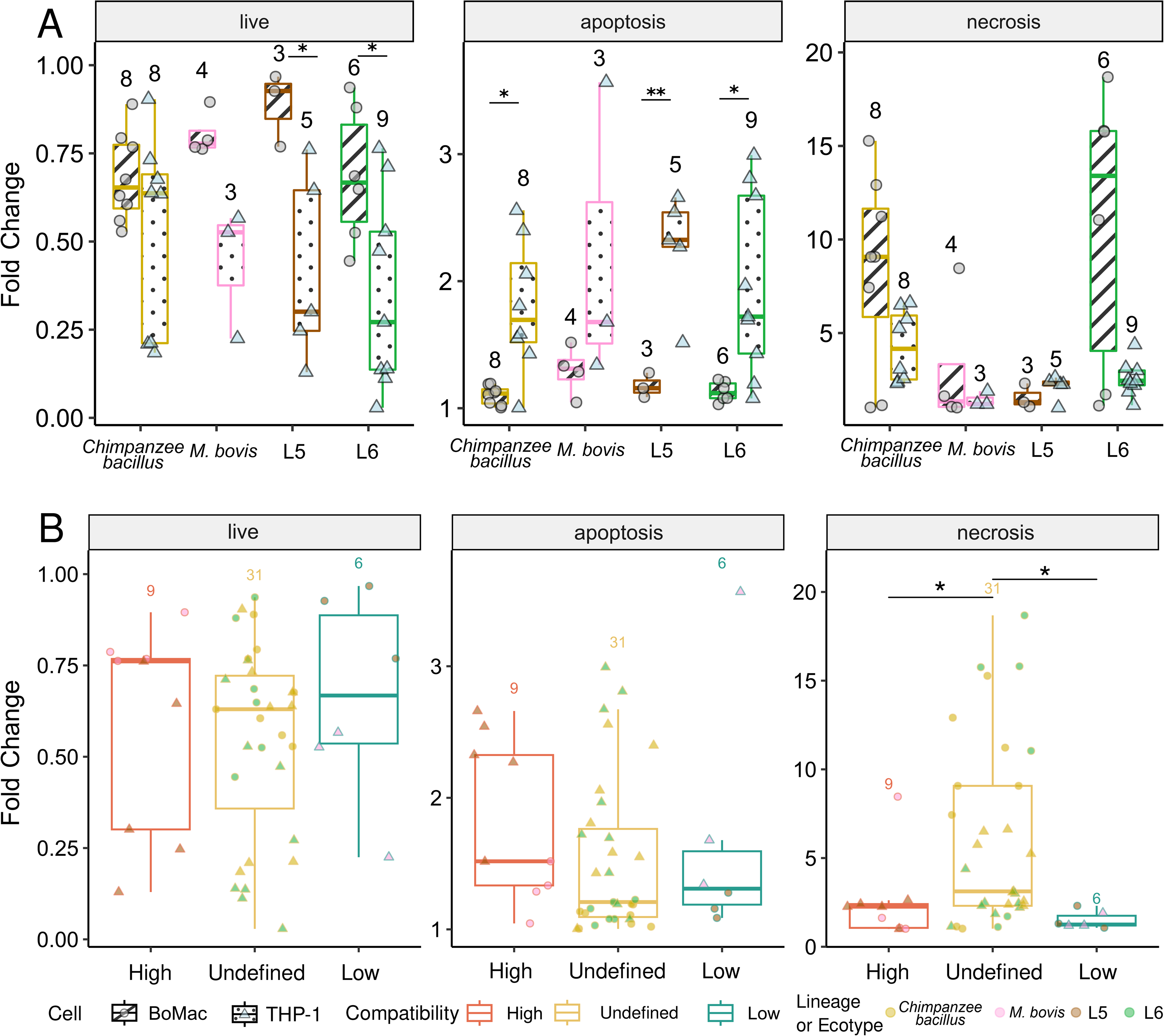
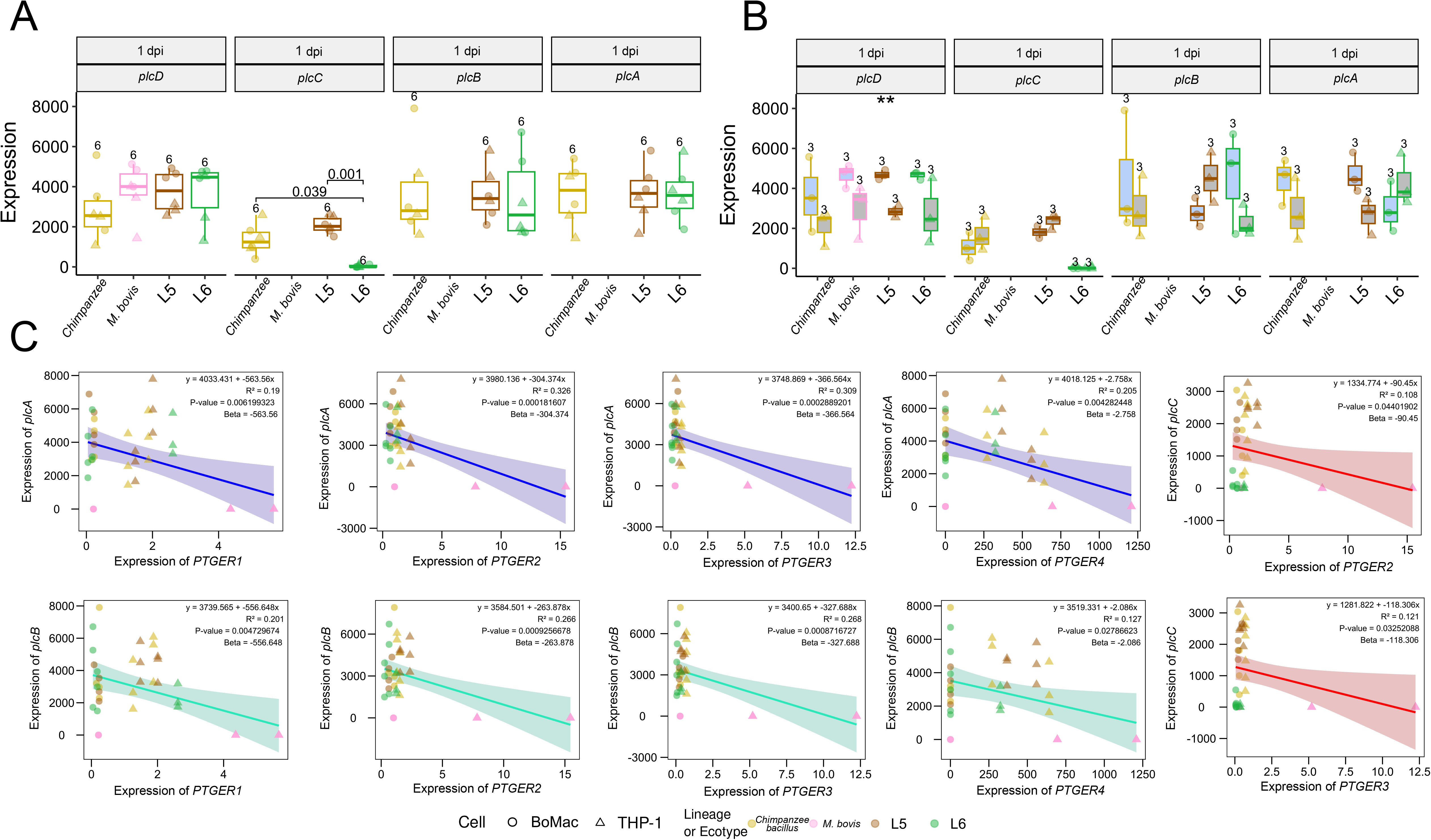

## Notes

### Competing Interest Statement

The authors have declared no competing interest.

### Summary of Updates

We added new results: Correlation between the signature virulence genes of M. tuberculosis and host receptors

## References

[1] Speranza E. Understanding virus-host interactions in tissues. Nat Microbiol. 2023;8:1397–1407.

[2] Finlay BB, McFadden G. Anti-Immunology: Evasion of the Host Immune System by Bacterial and Viral Pathogens. Cell. 2006;124:767–782.

[3] Rodriguez-Frandsen A, Alfonso R, Nieto A. Influenza virus polymerase: Functions on host range, inhibition of cellular response to infection and pathogenicity. Virus Res. 2015;209:23–38.

[4] Muylaert RL, Wilkinson DA, Kingston T, et al. Using drivers and transmission pathways to identify SARS-like coronavirus spillover risk hotspots. Nat Commun. 2023;14:6854.

[5] Tan CCS, Lam SD, Richard D, et al. Transmission of SARS-CoV-2 from humans to animals and potential host adaptation. Nat Commun. 2022;13:2988.

[6] Wattam AR, Foster JT, Mane SP, et al. Comparative phylogenomics and evolution of the Brucellae reveal a path to virulence. J Bacteriol. 2014;196:920–930.

[7] Gagneux S. Ecology and evolution of Mycobacterium tuberculosis. Nat Rev Microbiol. 2018;16:202–213.

[8] WHO. Global Tuberculosis Report 2023 [Internet]. [cited 2024 May 6]. Available from: https://www.who.int/teams/global-tuberculosis-programme/tb-reports/global-tuberculosis-report-2023.

[9] Brites D, Loiseau C, Menardo F, et al. A New Phylogenetic Framework for the Animal-Adapted Mycobacterium tuberculosis Complex. Front Microbiol [Internet]. 2018 [cited 2022 Nov 27];9. Available from: https://www.frontiersin.org/articles/10.3389/fmicb.2018.02820.

[10] Guyeux C, Senelle G, Le Meur A, et al. Newly Identified Mycobacterium africanum Lineage 10, Central Africa. Emerg Infect Dis. 2024;30:560–563.

[11] Coscolla M, Gagneux S, Menardo F, et al. Phylogenomics of Mycobacterium africanum reveals a new lineage and a complex evolutionary history. Microb Genomics. 2021;7:000477.

[12] Ngabonziza JCS, Loiseau C, Marceau M, et al. A sister lineage of the Mycobacterium tuberculosis complex discovered in the African Great Lakes region. Nat Commun. 2020;11:2917.

[13] Brites D, Gagneux S. Co-evolution of Mycobacterium tuberculosis and Homo sapiens. Immunol Rev. 2015;264:6–24.

[14] Coscolla M, Gagneux S. Consequences of genomic diversity in Mycobacterium tuberculosis. Semin Immunol. 2014;26:431–444.

[15] Berg S, Smith NH. Why doesn’t bovine tuberculosis transmit between humans? Trends Microbiol. 2014;22:552–553.

[16] Evans JT, Smith EG, Banerjee A, et al. Cluster of human tuberculosis caused by Mycobacterium bovis: evidence for person-to-person transmission in the UK. The Lancet. 2007;369:1270–1276.

[17] Ghielmetti G, Coscolla M, Ruetten M, et al. Tuberculosis in Swiss captive Asian elephants: microevolution of Mycobacterium tuberculosis characterized by multilocus variable-number tandem-repeat analysis and whole-genome sequencing. Sci Rep. 2017;7:14647.

[18] Whelan AO, Coad M, Cockle PJ, et al. Revisiting Host Preference in the Mycobacterium tuberculosis Complex: Experimental Infection Shows M. tuberculosis H37Rv to Be Avirulent in Cattle. PLOS ONE. 2010;5:e8527.

[19] Villarreal-Ramos B, Berg S, Whelan A, et al. Experimental infection of cattle with Mycobacterium tuberculosis isolates shows the attenuation of the human tubercle bacillus for cattle. Sci Rep. 2018;8:894.

[20] Yeboah-Manu D, Asare P, Asante-Poku A, et al. Spatio-Temporal Distribution of Mycobacterium tuberculosis Complex Strains in Ghana. PloS One. 2016;11:e0161892.

[21] Otchere ID, Coscollá M, Sánchez-Busó L, et al. Comparative genomics of Mycobacterium africanum Lineage 5 and Lineage 6 from Ghana suggests distinct ecological niches. Sci Rep. 2018;8:11269.

[22] Coscolla M, Lewin A, Metzger S, et al. Novel Mycobacterium tuberculosis Complex Isolate from a Wild Chimpanzee. Emerg Infect Dis. 2013;19:969–976.

[23] Ates LS. New insights into the mycobacterial PE and PPE proteins provide a framework for future research. Mol Microbiol. 2020;113:4–21.

[24] Mengaud J, Braun-Breton C, Cossart P. Identification of phosphatidylinositol-specific phospholipase C activity in Listeria monocytogenes: a novel type of virulence factor? Mol Microbiol. 1991;5:367–372.

[25] Terada LS, Johansen KA, Nowbar S, et al. Pseudomonas aeruginosa hemolytic phospholipase C suppresses neutrophil respiratory burst activity. Infect Immun. 1999;67:2371–2376.

[26] Assis PA, Espíndola MS, Paula-Silva FW, et al. Mycobacterium tuberculosis expressing phospholipase C subverts PGE2 synthesis and induces necrosis in alveolar macrophages. BMC Microbiol. 2014;14:128.

[27] Kaul V, Bhattacharya D, Singh Y, et al. An Important Role of Prostanoid Receptor EP2 in Host Resistance to Mycobacterium tuberculosis Infection in Mice. J Infect Dis. 2012;206:1816–1825.

[28] Borrell S, Trauner A, Brites D, et al. Reference set of Mycobacterium tuberculosis clinical strains: A tool for research and product development. PLOS ONE. 2019;14:e0214088.

[29] Goig GA, Cancino-Muñoz I, Torres-Puente M, et al. Whole-genome sequencing of Mycobacterium tuberculosis directly from clinical samples for high-resolution genomic epidemiology and drug resistance surveillance: an observational study. Lancet Microbe. 2020;1:e175–e183.

[30] BBMap: A Fast, Accurate, Splice-Aware Aligner. United States. Department of Energy. Office of Science; 2014.

[31] Andrew S. FastQC: a quality control tool for high throughput sequence data [Internet]. 2010 [cited 2022 Nov 15]. Available from: https://www.bioinformatics.babraham.ac.uk/projects/fastqc/.

[32] Dobin A, Davis CA, Schlesinger F, et al. STAR: ultrafast universal RNA-seq aligner. Bioinforma Oxf Engl. 2013;29:15–21.

[33] Liao Y, Smyth GK, Shi W. The R package Rsubread is easier, faster, cheaper and better for alignment and quantification of RNA sequencing reads. Nucleic Acids Res. 2019;47:e47.

[34] Rau A, Gallopin M, Celeux G, et al. Data-based filtering for replicated high-throughput transcriptome sequencing experiments. Bioinformatics. 2013;29:2146–2152.

[35] Love MI, Huber W, Anders S. Moderated estimation of fold change and dispersion for RNA-seq data with DESeq2. Genome Biol. 2014;15:550.

[36] Durinck S, Spellman PT, Birney E, et al. Mapping identifiers for the integration of genomic datasets with the R/Bioconductor package biomaRt. Nat Protoc. 2009;4:1184–1191.

[37] Wu T, Hu E, Xu S, et al. clusterProfiler 4.0: A universal enrichment tool for interpreting omics data. Innov Camb Mass. 2021;2:100141.

[38] Sayols S. rrvgo: a Bioconductor package for interpreting lists of Gene Ontology terms. MicroPublication Biol. 2023:10.17912/micropub.biology.000811.

[39] Putri GH, Anders S, Pyl PT, et al. Analysing high-throughput sequencing data in Python with HTSeq 2.0. Bioinformatics. 2022;38:2943–2945.

[40] Anders S, Reyes A, Huber W. Detecting differential usage of exons from RNA-seq data. Genome Res. 2012;22:2008–2017.

[41] Koo M-S, Subbian S, Kaplan G. Strain specific transcriptional response in Mycobacterium tuberculosis infected macrophages. Cell Commun Signal CCS. 2012;10:2.

[42] Leisching G, Pietersen R-D, van Heerden C, et al. RNAseq reveals hypervirulence-specific host responses to M. tuberculosis infection. Virulence. 2017;8:848–858.

[43] Zhou S, Goldstein S, Place M, et al. A clone-free, single molecule map of the domestic cow (Bos taurus) genome. BMC Genomics. 2015;16:644.

[44] Harding CV, Boom WH. Regulation of antigen presentation by Mycobacterium tuberculosis: a role for Toll-like receptors. Nat Rev Microbiol. 2010;8:296–307.

[45] Pu W, Zhao C, Wazir J, et al. Comparative transcriptomic analysis of THP-1-derived macrophages infected with Mycobacterium tuberculosis H37Rv, H37Ra and BCG. J Cell Mol Med. 2021;25:10504–10520.

[46] Kessler M, Zielecki J, Thieck O, et al. Chlamydia trachomatis disturbs epithelial tissue homeostasis in fallopian tubes via paracrine Wnt signaling. Am J Pathol. 2012;180:186–198.

[47] Pieters J. Mycobacterium tuberculosis and the macrophage: maintaining a balance. Cell Host Microbe. 2008;3:399–407.

[48] Amaral EP, Costa DL, Namasivayam S, et al. A major role for ferroptosis in Mycobacterium tuberculosis-induced cell death and tissue necrosis. J Exp Med. 2019;216:556–570.

[49] Gutierrez MG, Master SS, Singh SB, et al. Autophagy is a defense mechanism inhibiting BCG and Mycobacterium tuberculosis survival in infected macrophages. Cell. 2004;119:753–766.

[50] DesJardin LE, Kaufman TM, Potts B, et al. Mycobacterium tuberculosis-infected human macrophages exhibit enhanced cellular adhesion with increased expression of LFA-1 and ICAM-1 and reduced expression and/or function of complement receptors, FcgammaRII and the mannose receptor. Microbiol Read Engl. 2002;148:3161–3171.

[51] Marrero J, Trujillo C, Rhee KY, et al. Glucose Phosphorylation Is Required for Mycobacterium tuberculosis Persistence in Mice. PLOS Pathog. 2013;9:e1003116.

[52] Pellegrino E, Aylan B, Bussi C, et al. Peroxisomal ROS control cytosolic Mycobacterium tuberculosis replication in human macrophages. J Cell Biol. 2023;222:e202303066.

[53] Lerner TR, Borel S, Gutierrez MG. The innate immune response in human tuberculosis. Cell Microbiol. 2015;17:1277–1285.

[54] Kalam H, Fontana MF, Kumar D. Alternate splicing of transcripts shape macrophage response to Mycobacterium tuberculosis infection. PLOS Pathog. 2017;13:e1006236.

[55] Mathy NL, Scheuer W, Lanzendörfer M, et al. Interleukin-16 stimulates the expression and production of pro-inflammatory cytokines by human monocytes. Immunology. 2000;100:63–69.

[56] Ates LS, Sayes F, Frigui W, et al. RD5-mediated lack of PE_PGRS and PPE-MPTR export in BCG vaccine strains results in strong reduction of antigenic repertoire but little impact on protection. PLoS Pathog. 2018;14:e1007139.

[57] Sheppe AEF, Edelmann MJ. Roles of Eicosanoids in Regulating Inflammation and Neutrophil Migration as an Innate Host Response to Bacterial Infections. Infect Immun. 2021;89:e0009521.

[58] Gagneux S, DeRiemer K, Van T, et al. Variable host–pathogen compatibility in Mycobacterium tuberculosis. Proc Natl Acad Sci. 2006;103:2869–2873.

[59] Hiza H, Zwyer M, Hella J, et al. Bacterial diversity dominates variable macrophage responses of tuberculosis patients in Tanzania. Sci Rep. 2024;14:9287.

[60] Maqsood MI, Matin MM, Bahrami AR, et al. Immortality of cell lines: challenges and advantages of establishment. Cell Biol Int. 2013;37:1038–1045.

