## Supplemental material for "Host-strain compatibility influences transcriptional responses in *Mycobacterium tuberculosis* infections"

#### Figures

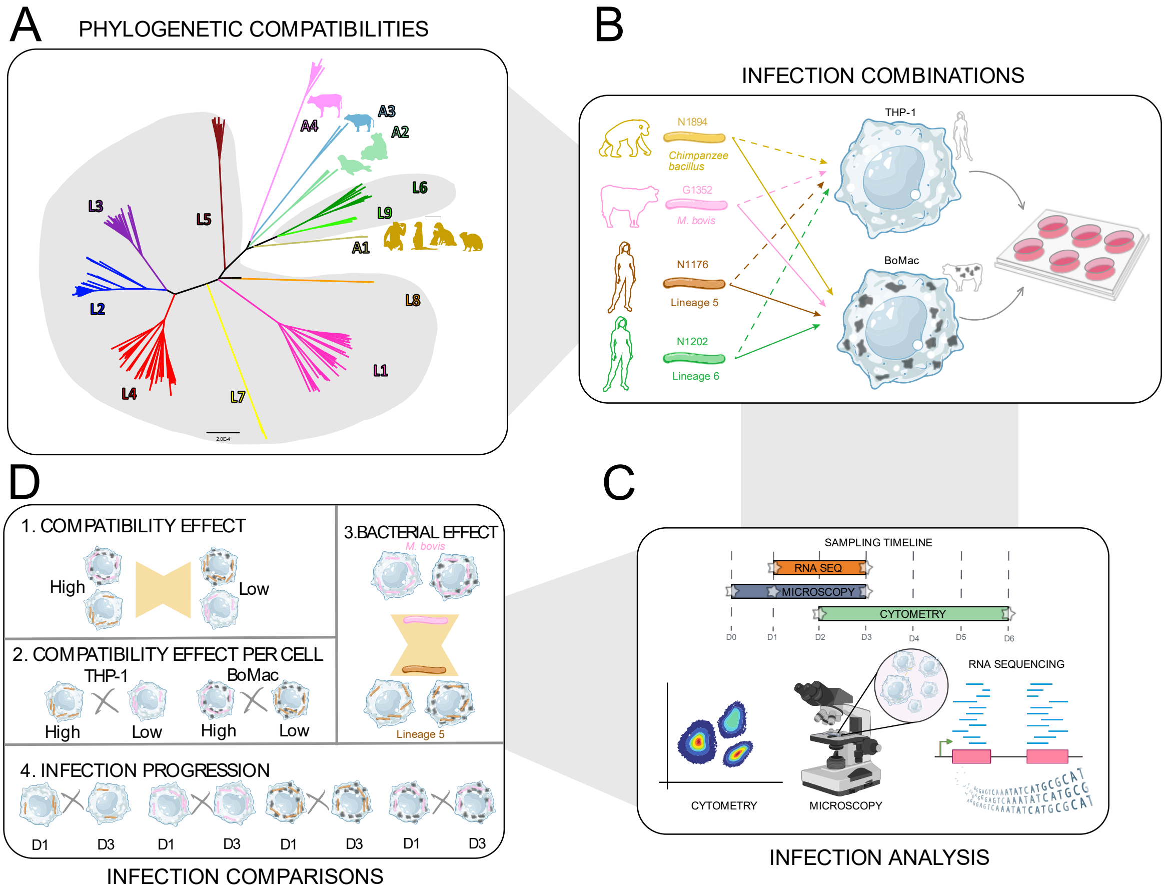


**Figure S1. General experimental setup of crossed-infections in *M. tuberculosis*.**
**A)** Phylogenetic relationships among 13 *M. tuberculosis* lineages, with animal-associated lineages labelled with 'A' and a silhouette of their most common hosts, and human host-associated lineages with 'L' and shaded in grey. **B)** Representation of the crossed-infection experiments using four *M. tuberculosis* strains on THP-1 and BoMac cell lines. **C)** Infections setup times and their corresponding analysis techniques including cytometry, microscopy, RNA extraction, and sequencing. **D)** Transcriptional comparisons for all infection combinations (strain-cell), for the comparisons (1-4) we have concatenated and split analysis for 1-day post-infection (dpi) and 3 dpi.

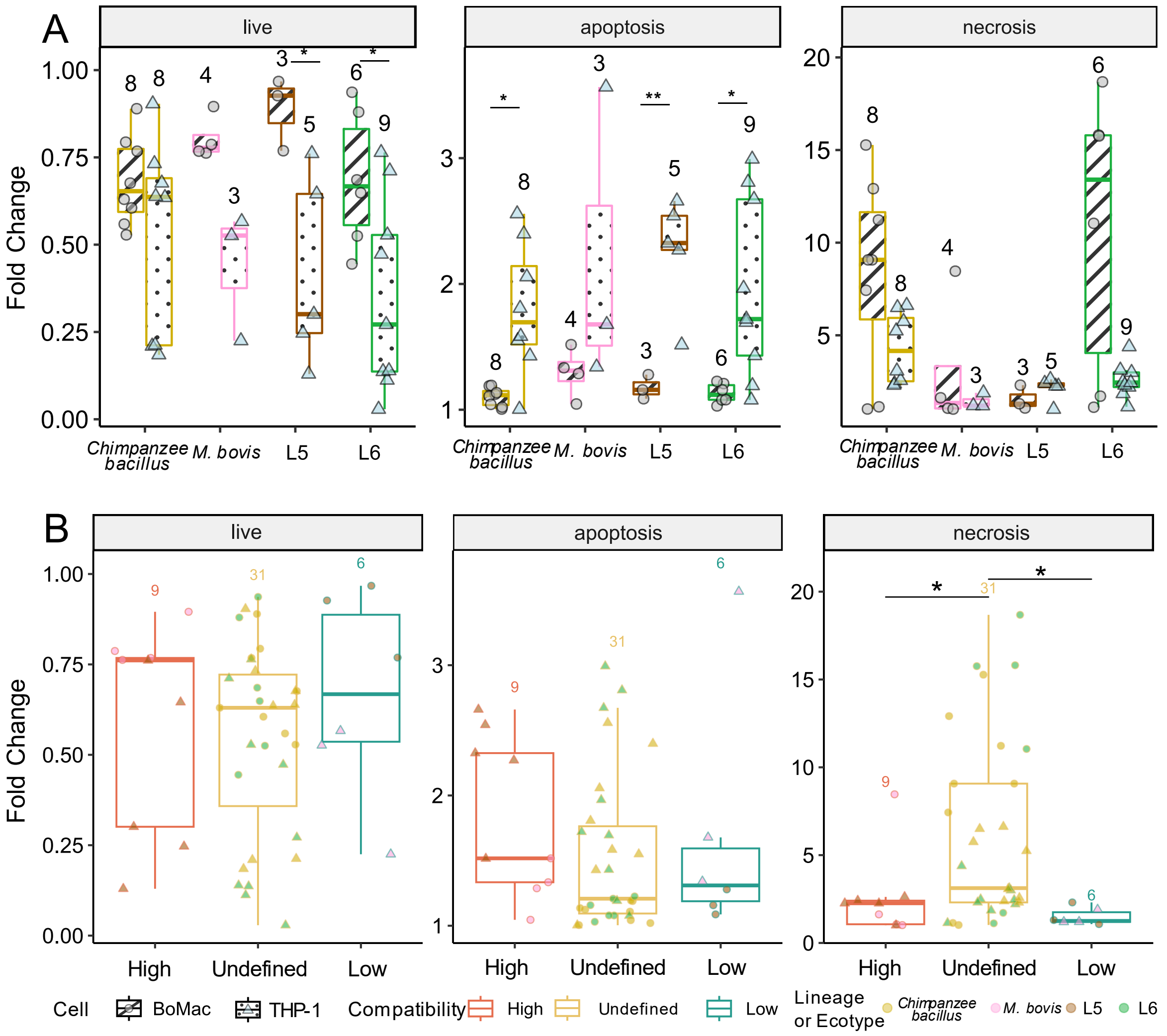


**Figure S2. Cell death assessed by cytometry in cross-infections of THP-1 and BoMac cells for 2- and 6-days post-infection. A)** Fold change of each cell death for THP-1 and BoMac cell infections, evaluated by T-test. **B)** Fold change of cell death for each compatibility group (high, undefined, low). Fold change was calculated by comparing sample values to the control mean. Significant differences from both statistical tests are shown as : $p \leq 0.05$; *: $p \leq 0.01$; *: $p \leq 0.001$; **: $p \leq 0.0001$.

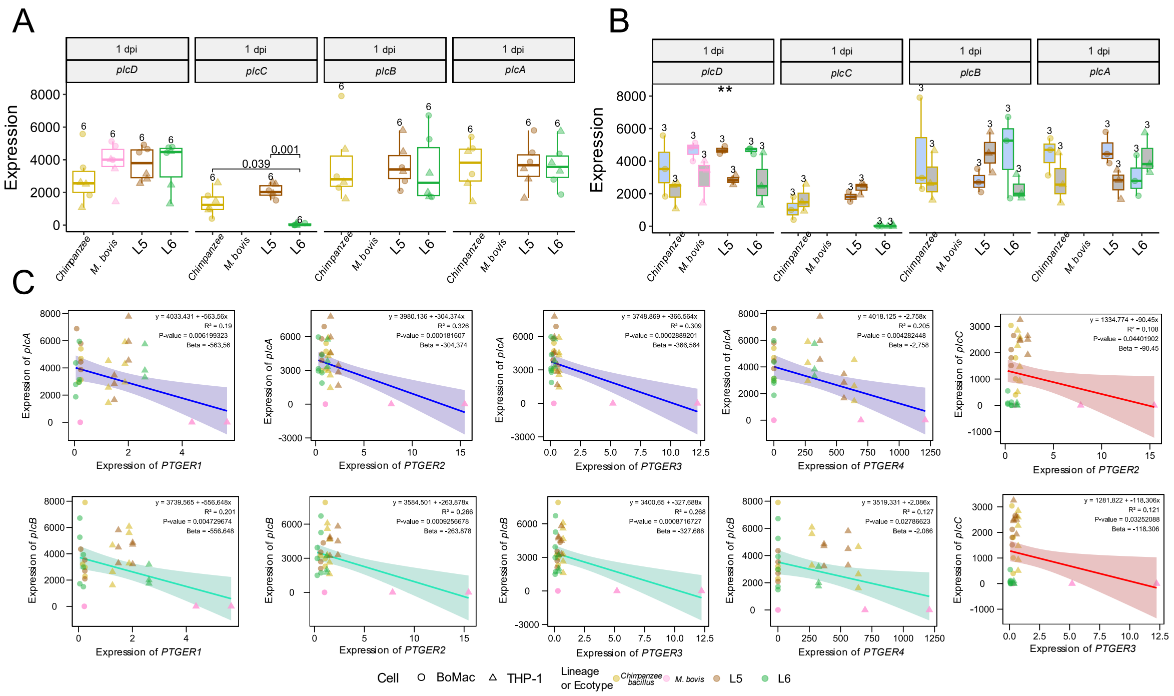

**Figure S3. Expression phospholipases C and *PTGER* genes in BoMac and THP-1 infections at 1 dpi. A)** Expression of *plc*s genes from *M. tuberculosis* strains in THP-1 and BoMac infections at 1 dpi. Only significant results are shown for clarity (*: p ≤ 0.05; **: p ≤ 0.01; ***: p ≤ 0.001; ****: p ≤ 0.0001). **B)** Comparison of *plc*s expression in THP-1 and BoMac cell infections at 1 dpi, evaluated by the T-test. **C)** Correlations between the expression of PTGERs and *plcA* (blue line), *plcB* (green line) and *plcC* (red line).

### Tables (Pending release)

**Table S1.** Infection ratio per sample.
**Table S2.** Percentage of cell death per sample.
**Table S3.** Accession numbers of sequencing raw reads per sample.
**Table S4.** Expression of phospholipases c (A, B, C, and D) per sample.
**Table S5.** Primers to quantify the expression of phospholipases c of *M. tuberculosis* strains.
**Table S6.** Differentially expressed orthologous genes in the comparison of high vs. low compatibilities in both BoMac and THP-1 cell lines. The comparison groups are: genes upregulated in high compatibility compared to low compatibility (High_orthologs), genes upregulated in low compatibility compared to high compatibility (Low_orthologs); Genes upregulated in L5 strain compared to *M. bovis* strain (L5_orthologs), and genes upregulated in *M. bovis* strains compared to L5 strain (Mbovis_orthologs). Only significant results are represented (p adj. < 0.05)
**Table S7.** Gene enrichment results for the differentially expressed orthologous genes (DEOGs) across all gene ontologies and KEGG pathways in the comparison of high vs. low compatibilities in both BoMac and THP-1 cell lines. The comparison groups are: genes upregulated in high compatibility compared to low compatibility (High_orthologs), genes upregulated in low compatibility compared to high compatibility (Low_orthologs), genes upregulated in L5 strain compared to *M. bovis* strain (L5_orthologs), and genes upregulated in *M. bovis* strains compared to L5 strain (Mbovis_orthologs). Only significant results are represented (p adj. < 0.05).
**Table S8.** Differentially expressed genes in the comparison of high vs. low compatibilities in both BoMac (High_Low_BoMac) and THP-1 (High_Low_THP-1) cell lines.
**Table S9.** Gene enrichment results for the differentially expressed genes (DEGs) across all gene ontologies and KEGG pathways in the comparison of high vs. low compatibilities in both BoMac (BoMac_High or BoMac_Low) and THP-1 (THP-1_High or THP-1_Low) cell lines. Only significant results are represented (p adj. < 0.05).
**Table S10.** Differentially expressed genes in the comparison of 1 dpi vs. 3 dpi in high (High_1dpi or High_3dpi) and low (Low_1dpi or Low_3dpi) compatibility. Only significant results are represented (p adj. < 0.05)
**Table S11.** Gene enrichment results for the differentially expressed genes (DEGs) across all gene ontologies and KEGG pathways in the comparison of 1 dpi vs. 3 dpi in high (High_1dpi or High_3dpi) and low (Low_1dpi or Low_3dpi) compatibilities. Only significant results are represented (p adj. < 0.05).
**Table S12.** Correlation results between gene expression of DEOGs and microscopy ratio.
**Table S13.** Differentially exon usage in the comparison high vs. low compatibility in THP-1 cells, genes harbouring differentially exons usages are labelled as genes with exons associated to high and compatibility (L5_and_Mbovis), only high compatibility (L5) and only low (Mbovis) compatibility. Only significant results are represented (p adj. < 0.05).
**Table S14.** Gene enrichment results for the differentially exon usage (DEUs) across all gene ontologies and KEGG pathways in the comparison of high (L5) vs. low (Mbovis) compatibility in THP-1. Only significant results are represented (p adj. < 0.05).
